# MetaProFi: A protein-based Bloom filter for storing and querying sequence data for accurate identification of functionally relevant genetic variants

**DOI:** 10.1101/2021.08.12.456081

**Authors:** Sanjay K. Srikakulam, Sebastian Keller, Fawaz Dabbaghie, Robert Bals, Olga V. Kalinina

**Author notes:** **Corresponding author:** Prof. Dr. Olga V. Kalinina, Helmholtz Institute for Pharmaceutical Research Saarland (HIPS), Helmholtz Centre for Infection Research (HZI), Campus E8.1, 66123 Saarbrücken, Germany and Medical Faculty, Saarland University, 66421 Homburg, Germany, Campus E8.1, 66123 Saarbrücken, Germany.

## Abstract

Technological advances of next-generation sequencing present new computational challenges to develop methods to store and query these data in time- and memory-efficient ways. We present MetaProFi (https://github.com/kalininalab/metaprofi), a Bloom filter-based tool that, in addition to supporting nucleotide sequences, can for the first time directly store and query amino acid sequences and translated nucleotide sequences, thus bringing sequence comparison to a more biologically relevant protein level. Owing to the properties of Bloom filters, it has a zero false-negative rate, allows for exact and inexact searches, and leverages disk storage and Zstandard compression to achieve high time and space efficiency. We demonstrate the utility of MetaProFi by indexing UniProtKB datasets at organism- and at sequence-level in addition to the indexing of Tara Oceans dataset and the 2585 human RNA-seq experiments, showing that MetaProFi consumes far less disk space than state-of-the-art-tools while also improving performance.

## Introduction

With the technological advancement in the field of next-generation sequencing (NGS) over the past decade, there is a rapid growth in the amount of available biological sequencing data in public databases e.g., European Nucleotide Archive (ENA)^[1]^, and Sequence Read Archive (SRA)^[2]^. NGS data has become an invaluable resource in various fields of life science research. As the size of the databases reached petabyte-scale it has become difficult to support online searches in these databases. As great power comes with great responsibility, the requirement for tools to process, store and query large collections of data without high memory and storage requirements constitutes a computational challenge. Analyzing these abundant data will lead to opportunities for great scientific discoveries.

To reduce the required time and memory, a probabilistic data structure can be used to summarize and deliver fast approximate answers. Probabilistic membership data structures aid in determining whether an item/element is present or not. Several such data structures are already available, e.g. Bloom filter, Cuckoo filter, Quotient filter, Count-min sketch, HyperLogLog, MinHash, etc. These data structures differ in their ability to offer approximation or definite answers. Many of these data structures are predominantly used in streaming applications and database lookups before performing any expensive operations. For example, before fetching data from a database the key lookups are made to check for data existence. These data structures guarantee zero false negatives, thus if such a data structure returns an answer of non-existent key, then the database fetch/read is not required, saving time.

In the last few years several tools (SBT^[3]^, Mantis^[4]^, Howde-SBT^[5]^, BIGSI^[6]^, REINDEER^[7]^, kmtricks^[8]^) that make use of these probabilistic data structures (and their variants) have been made available to build custom indexes since tools like BLAST^[9]^ do not scale to modern-day large collections of data. The custom indexes are built either of the raw sequencing data or of the data from the curated databases and later allow the sequences of interest to be queried against these custom indexes to find which samples in the index contain the query sequence. Although several tools are available, none of them supports amino acid sequence indexing, creating a major gap in the field.

To this end, we introduce MetaProFi, a first of its kind tool for indexing amino acid sequences that also supports nucleotide sequence (canonical) indexing, using Bloom filters as the underlying data structure. Bloom filter (Figure 1) is a probabilistic set-membership data structure that stores the presence or absence of items/elements in a bit-vector (an array of binary values) and can be queried for presence or absence. Bloom filters guarantee zero false negatives. The bit-vector is filled with binary values where a zero indicates absence and one indicates presence: we start by filling the bit-vector with zeros; for each given string, we apply *h* hash functions and use the return value of hashing as an index in the bit-vector to flip the zero in the corresponding index position to one. Collisions can be avoided by using large Bloom filters and perfect hash functions, but in a realistic setting, this is not possible, thus false positives arise. Their number can be reduced by increasing the size of the Bloom filter and manipulating the number of hash functions used. An alternative to Bloom filters can be Quotient or Cuckoo filters, but their runtime degrades as the data structure is being filled up, and they are more demanding in terms of storage and memory, which makes them difficult to scale to millions of samples or datasets.

**Figure 1:**
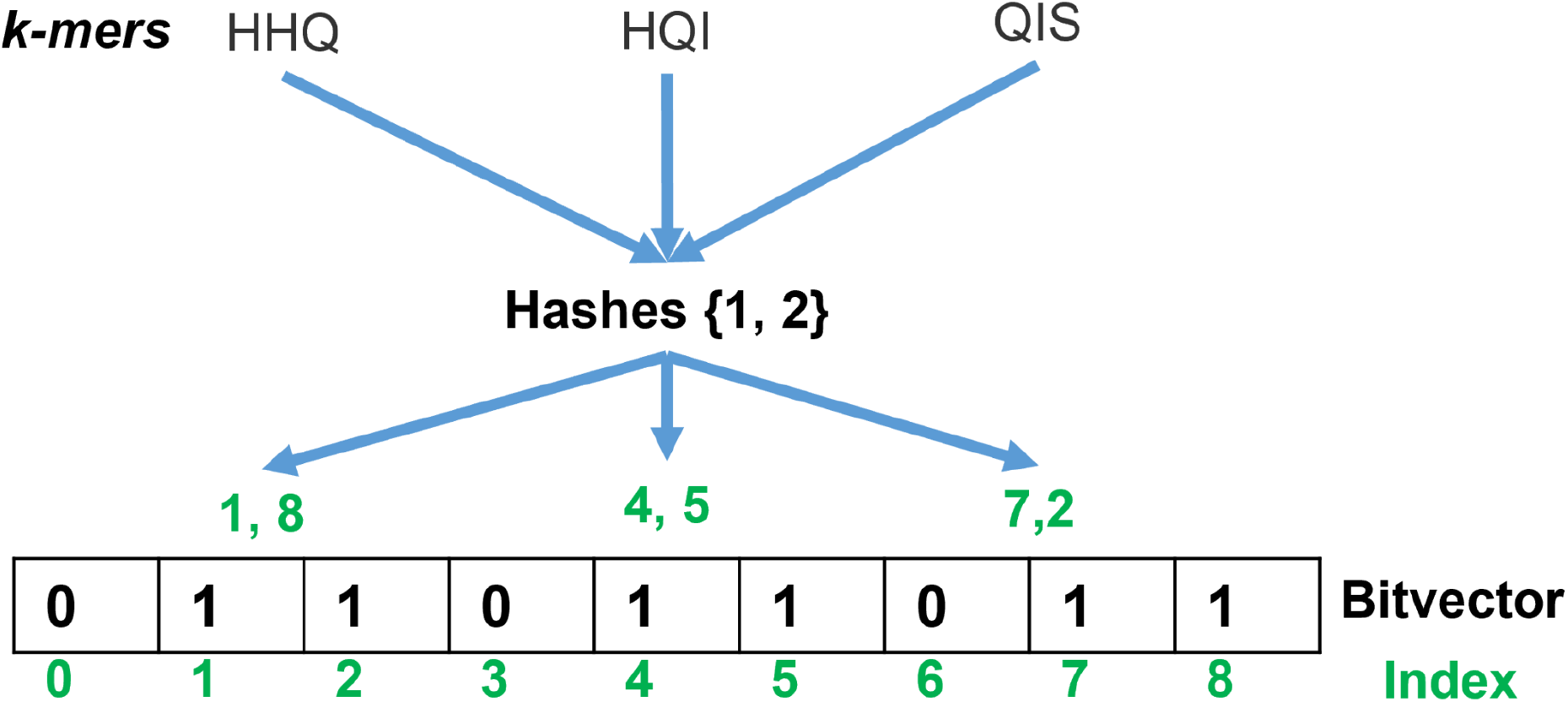
Bloom filter data structure: Two hash functions are applied to each of the three strings and the return values of hashing are used as an index in the bit-vector to flip the zero in the corresponding index position to one.

MetaProFi combines the power of a variant of Bloom filter data structure which we call packed Bloom filter (see Methods for details) with chunked disk storage and Zstandard compression algorithm to construct the Bloom filter matrix efficiently to index all the observed *k*-mers (presence/absence) for fast queries. We demonstrate superior performance compared to previous studies. MetaProFi enables querying assembled sequence contigs from metagenomics data against a custom amino acid database. MetaProFi serves for various applications: one can index all the sequences available in UniProtKB^[10]^ which allows for fast alignment-free sequence search or to find resistance-associated genes by indexing resistance-associated markers. MetaProFi does not only perform exact sequence searches but also allows one to do an approximate sequence search by providing a threshold (*T*). As a proof of concept, we have constructed a MetaProFi index for UniProtKB bacterial sequences (amino acids) in two different levels (organism-level and sequence-level), for the Tara Oceans^[11]^ (nucleotides) dataset, and for the 2585 human RNA-seq experiments comprising blood, brain and breast samples to demonstrate the storage, memory, run time, scalability and query performance.

## Results

MetaProFi allows the indexing of large numbers of samples/datasets. MetaProFi (Figure 2) combines the power of a probabilistic data structure with compression algorithms and chunked storage to store large Bloom filters and to create indexes for fast querying. MetaProFi constructs a Bloom filter matrix (Figure 2a) for the given samples, by (1) reading each sequence, (2) extracting *k*-mers from the sequence (3) applies *h* hash functions, resulting in *h* integers (one integer per hash function), (4) flipping the respective bits in the Bloom filter to indicate that the *k*-mer is present. This is repeated for all sequences in all samples and a two-dimensional (2D) Bloom filter matrix is constructed on disk where each column represents a Bloom filter of a sample. MetaProFi then reads multi-row chunks from the 2D Bloom filter matrix (see methods for details) without consuming too much memory and creates an index (Figure 2b) that can be utilized for fast querying. MetaProFi allows adding new samples to the index by creating a Bloom filter matrix for the new dataset (set of samples or FASTA/FASTQ files) and then appending the new data (columns) to the end of the existing index.

**Figure 2:**
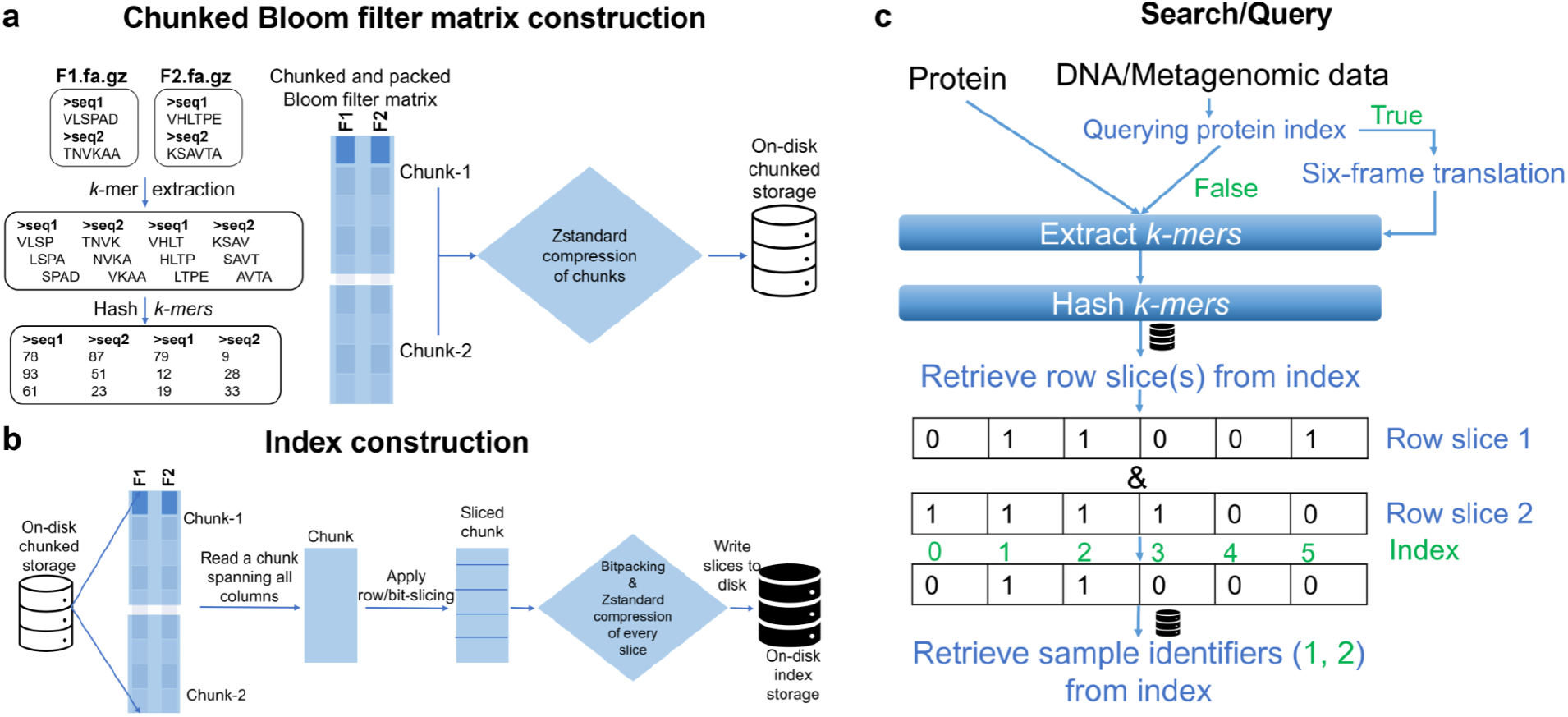
MetaProFi pipeline: (a) Chunked Bloom filter matrix construction, (b) Index construction, and (c) Search/query pipeline

To query the index (Figure 2c) MetaProFi extracts all the *k*-mers in a given sequence and then applies *h* hash functions and acquires *h* integers. At this point, one can consider the MetaProFi index as a hash table, and MetaProFi extracts the row/bit-slice from this hash table using the *h* integers and performs bitwise operations to find sample(s) containing the query. MetaProFi allows exact query search expecting every *k*-mer in the query sequence to be present, and also supports approximate search using a threshold (*T*) when only the fraction of *k*-mers larger than *T* have to be found.

## MetaProFi performance

To evaluate the performance of MetaProFi, we used UniProtKB, Tara Oceans and 2585 RNA-seq datasets for the index construction. For UniProtKB, two types of indexes were created: one at the organism level and the other at the sequence level (See Methods for details). Since no other tool is available to perform amino acid *k*-mer indexing we added support for nucleotide indexing to MetaProFi to enable comparison against other tools and we indexed the Tara Oceans and the RNA-seq datasets for benchmarking purposes.

For UniProtKB organism-level indexing, the dataset was constructed by extracting individual bacterial sequences using their accession ids with criteria on the minimum length of the sequence (*k* = 11) and then grouping them by their organism name (OS field value in the fasta header) to obtain 100,384 uncompressed individual fasta files containing a total of 46,511,863,142 *k*-mers. MetaProFi constructs the Bloom filter matrix in 38.05 min and the index in under 1290.59 min using under 60 GiB of RAM and under 135 GiB of disk space (Table 1). Storage size is directly proportional to the size of the Bloom filter and number of samples in the dataset: if MetaProFi had used a regular Bloom filter, it would require 6.85 TiB (size of the Bloom filter times number of Fasta files/samples) disk space for storing the uncompressed Bloom filter matrix. With MetaProFi’s optimizations and techniques, we provide a 50-fold compression and can construct Bloom filter matrices for a large number of datasets or use very large Bloom filters that have a low false-positive rate, while still storing them efficiently.

**Table 1:** MetaProFi organism-level indexing results for UniProtKB bacterial dataset

| Organism-level | Time (min) | RAM (GiB) | Disk (GiB) |
| --- | --- | --- | --- |
| Bloom filter matrix | 38.05 | <60 | 139 |
| MetaProFi index | 1290.59 | <60 | 135 |

To demonstrate MetaProFi’s scalability we used all the bacterial sequences that were extracted from the UniProtKB dataset, 334,984 sequences from Swiss-Prot and 151,450,171 sequences from TrEMBL. These were stored as one compressed file (referred to as UniProtKB bacterial dataset). MetaProFi supports sequence-level indexing with one caveat that a dedicated file index (see Methods for details) will be constructed for the input file and this may require a maximum of half of the original file’s storage. To demonstrate this, we constructed the minimalistic index for the compressed UniProtKB bacterial dataset of size 32 GiB. MetaProFi’s LMDB (Lightning Memory-Mapped Database) (https://lmdb.readthedocs.io/en/release/) indexer took 17 minutes to construct the index and used 11 GiB of storage and a maximum of 1 GiB RAM. Using LMDB as the underlying database reduces the RAM consumption as we do not retain the index in memory and write the data for every sequence to the disk as soon as they are populated. The 1 GiB of RAM consumption is due to the Gzip index construction for Gzip compressed input files and as soon as it is written to the disk, the memory is freed and thus more most of the time only a few MiB of RAM for constructing the FASTA/FASTQ sequence index are required. Using the LMDB index of the UniProtKB bacterial dataset (151,785,155 sequences) we created a sequence-level MetaProFi index. The Bloom filter matrix and index construction time are comparable with the organism-level case. The disk requirements are twice as large, and RAM consumption is comparable (Table 2). These results show that MetaProFi is scalable to hundreds of millions of samples.

**Table 2:** MetaProFi sequence-level indexing results for UniProtKB bacterial dataset

| Sequence-level | Time (min) | RAM (GiB) | Disk (GiB) |
| --- | --- | --- | --- |
| Bloom filter matrix | 66.33 | <60 | 232 |
| MetaProFi index | 862.57 | <60 | 210 |

To compare MetaProFi with the state of the art tools, we used kmtricks^[8]^, a *k*-mer counting tool that allows building Bloom filters that can be used for constructing a *k*-mer index using the HowDe-SBT variant implemented in its package. Though other tools like BIGSI, HowDe-SBT, and Mantis aim to index large collections of data, indexing Tara Oceans like dataset seems to be not feasible^[8]^. We built a MetaProFi index for the Tara Oceans dataset (Table 3, see Methods for details of filter construction) containing a total of 3,431,551,187,218 *k*-mers (*k* = 31). MetaProFi constructed the Bloom filter matrix in 2998.34 minutes and index under 252.24 minutes using less than 58 GiB RAM and 640 GiB of disk space and 70 GiB RAM and 751 GiB of disk space respectively (Table 3).

**Table 3:** Tara Oceans dataset benchmark comparisons of MetaProFi and kmtricks

| Tara Ocean dataset | MetaProFi |  |  |  | kmtricks + HowDe-Sbt |  |  |  |
| --- | --- | --- | --- | --- | --- | --- | --- | --- |
|  | Time (min) | RAM (GiB) | CPU cores | Disk (GiB) | Time (min) | RAM (GiB) | CPU cores | Disk (TiB) |
| Bloom filter matrix | 2998.34 | <58 | 64 | 640 | - | <50 | 64 | 4.8 |
| Index | 252.24 | 70 | 64 | 751 | - | - | - | - |

To demonstrate MetaProFi’s query performance, we randomly selected 1000 reads from the 495 FASTQ files of the Tara Oceans dataset used for constructing the MetaProFi index. These 1000 reads were queried against the Tara Oceans MetaProFi index using exact search (*T* = 100%) and approximate search (*T* = 40%) and query run times were 12.877 seconds and 13.048 seconds respectively.

To further compare MetaProFi’s performance with other tools such as SBT, Howde-SBT, Mantis, BIGSI and REINDEER, we indexed the 2585 human RNA-seq experiments data obtained from the SRA^[2]^. MetaProFi, unlike other tools, does not filter/remove *k*-mers, hence a total of 6,432,932,578,661 *k*-mers (*k* = 21) were present in the 2585 RNA-seq dataset. The performance metrics of the state-of-the-art tools were obtained from the literature (SBT^[3]^, Mantis^[4]^, Howde-SBT^[5]^, BIGSI^[6]^, REINDEER^[7]^) and compared to MetaProFi (Table 4). This comparison is an imprecise estimate of the actual performance, as it is not only architecture-dependent but also depends on the additional filtering techniques used. Nevertheless, MetaProFi demonstrates a clear advantage in terms of total disk usage and, most importantly, execution time: it can construct the index for the 2585 human RNA-seq samples in 240.168 minutes (Bloom filter matrix construction: 1399.15 minutes using 59 GiB RAM) and requires only 333 GiB disk space for storing 6,432,932,578,661 *k*-mers (*k* = 21) without employing any *k*-mer filtering techniques.

**Table 4:** Human RNA-seq (2585 samples) index construction resource requirements of MetaProFi and other tools as reported in the literature. N/A indicates that the value was not reported in the article.

| Tool | Total disk (GiB) | Index size (GiB) | Index build time (h) | RAM (GiB) | k-mer size |
| --- | --- | --- | --- | --- | --- |
| SBT | N/A | 200 | 34 | N/A | 20 |
| Howde-SBT | N/A | 15 | 9 | N/A | 20 |
| Mantis | 2546.57 | 32 | 16.5833 | 40 | 20 |
| BIGSI | N/A | 144 | N/A | N/A | 31 |
| REINDEER-presence/absence | 6800 | 36 | 40 | 27 | 21 |
| MetaProFi | 628 | 333 | 4.0028 | 56 | 21 |

We then used this index for querying 1000 transcripts (see Methods for details) using 20 CPU cores. MetaProFi required 349 seconds and 1.9 GiB (resident memory) RAM for an exact search (T=100%) and 359 seconds and 1.9 GiB (resident memory) RAM for an approximate search (T=75%).

Our benchmarking results show that MetaProFi reduces the amount of disk usage even for very large Bloom filters, compared to the state-of-the-art tools and constructs index in little time.

## Discussion

We developed MetaProFi, a first of its kind amino acid *k*-mer indexing tool with added support for indexing nucleotide sequences that efficiently indexes from tens of samples to hundreds of millions of samples with reduced memory, storage and run time than its predecessors. We demonstrated this through our proof of concept index construction for multiple datasets that we can grow our index horizontally (samples) or vertically (Bloom filter size) and yet requires less amount of storage and memory without compromising on the performance both in terms of building and querying. MetaProFi can be further developed to support distributed computing infrastructure in addition to the current single system-specific deployment setup.

Benchmarking MetaProFi against state-of-the-art tools presented us with significant difficulties. MetaProFi was able to build *k*-mer indices for multiple datasets demonstrating it can scale in any direction; on the contrary, we failed to reproduce the reported results for other tools. For example, we attempted to construct kmtricks^[8]^ Bloom filter of the Tara Oceans dataset three times using two different sets of parameters (see methods for details). The first attempt was terminated after 9 hrs due to the consumption of the entire local disk space of 2.9 TiB while using the default parameters. In the second attempt, we applied the same parameters as in the first but switched the output to a large non-RAID NVMe NFS filesystem. This attempt was manually terminated after 100 hrs, during which we observed a peak usage of 4.8 TiB of storage. The third attempt, where we used the parameters reported in kmtricks raised a SIGSEGV error and was terminated after 11 hrs while consuming a total of 2.1 TiB of storage. Hence, we could compare MetaProFi’s performance only to the reported figures, bearing in mind the possible differences in the hardware and software. Nevertheless, MetaProFi showed state-of-the-art or better than state-of-the-art performance in terms of both speed and required resources.

The most important feature of MetaProFi, besides its storage and runtime efficiency, is that it builds its index for amino acid sequences in the default scenario. This approach was taken with the intended goal in mind: to most efficiently store and query sequence data coming from metagenome samples, primarily of bacterial metagenomes. Storing protein data makes the search more flexible and allows many advantages, while the only disadvantage is that the information in non-coding regions is lost. We consider this a little loss, since bacterial genomes contain comparatively little non-coding sequence, and a lot of important markers are detected on the protein level, e.g. markers of antibiotic resistance. Other examples of potential advantages include very fast and efficient searches with the *k*-mer presence threshold *T* = 100% can be conducted for closely related, but not identical DNA sequences that contain no missense mutations on the protein level. On the other hand, remote homologs can be detected in cases, when the sequence similarity on the DNA level drops, but is still detectable on the protein level.

## Methods

### MetaProFi Bloom filter matrix construction

MetaProFi is developed in Python and is parallelized to take advantage of all the available CPU cores on today’s modern computing systems in order to achieve the best performance. MetaProFi accepts both FASTA and FASTQ formats and uses pyfastx^[12]^ Python library for parsing the files efficiently. First, MetaProFi creates an empty 2-dimensional (2D) Bloom filter matrix with M rows and N columns shape, where rows represent hash indices and each column represents a Bloom filter for an individual sample. MetaProFi splits the number of input samples into N chunks of samples meticulously to make sure at all times MetaProFi do not consume more memory than what the user is set in the config file, and for each such chunk constructs a 2D Bloom filter sub-matrix using the POSIX shared memory. POSIX shared memory allows efficient inter-process communication while sharing a portion of memory and is the fastest available mechanism for passing the data between multiple processes on the same host system. Using POSIX shared memory enables us to offer a zero-copy solution (zero wastage of memory) by sharing the large 2D sub-matrix between multiple processes. The rows in the 2D sub-matrix represent hash indices and each column in the sub-matrix represents a Bloom filter of a sample in the chunk. MetaProFi dedicates 85% of RAM set in the config file for Bloom filter matrix construction and uses the rest of the RAM for IO and other operations, thus MetaProFi makes sure to never consume more memory.

The smallest addressable unit in modern architectures are byte-addressable, even the boolean data type in Python requires 1 byte to store a true or false value. Therefore using one byte for storing the presence or absence of a *k*-mer in Bloom filters would cause an increase in the memory and storage requirements eight times for any Bloom filter compared to most economical theoretically possible data storing. Hence MetaProFi uses an 8-bit unsigned integer (UINT8) data type to store bits for eight k-mers by applying bit manipulations to a UINT8 integer (a packed Bloom filter). MetaProFi utilizes this technique to optimize the storage and memory requirement, making it applicable to large datasets containing many samples.

Once all *k*-mers present in a single chunk of samples are hashed and the respective bits are flipped on the shared 2D Bloom filter sub-matrix, the sub-matrix is chunked horizontally (row-wise). Each chunk is then compressed using Zstandard compression algorithm for further reducing the storage requirements and these compressed chunks are written in parallel to the empty on disk 2D matrix MetaProFi created initially using Zarr^[13]^ Python library. Zarr library enables storing chunked, compressed, N-dimensional arrays. This is repeated for all N sample chunks.

### MetaProFi indexing

In addition to the Bloom filter construction for a large number of datasets, MetaProFi also offers efficient *k*-mer indexing for faster querying. For storing the index, MetaProFi first creates an empty one dimensional (1D) byte matrix of size M (size of the Bloom filter), where each row in the 1D matrix represents a row/bit-slice across all samples (columns) from the 2D Bloom filter matrix. MetaProFi re-creates a 2D POSIX shared memory matrix of size *m* rows and N samples, where N is equal to the total number of samples in the Bloom filter matrix and *m* is the size of the chunk of rows that were inserted during 2D Bloom filter matrix construction. MetaProFi reads *m* rows from each chunk spanning all N sample chunks, on the fly unpacks the UINT8 packed Bloom filter to individual bits, and writes the several unpacked bits of *m* rows and N samples to the POSIX shared memory matrix in parallel. MetaProFi then chunks these *m* rows into smaller chunks and distributes the chunks to multiple processes. In each process, MetaProFi takes a row/bit-slice that spans across all samples and applies bit-packing and Zstandard compression algorithms to reduce the storage requirements for the index. MetaProFi then writes these small compressed row/bit-slices in parallel to the empty 1D byte matrix created initially using the same Zarr library. MetaProFi repeats the procedure until all rows from the 2D Bloom filter matrix are indexed. MetaProFi also supports updates to the index for adding new Bloom filters using the same procedure. For appending new data to the index, during the bit-packing and compression phase of the new data, MetaProFi reads old data from the existing index and decompresses the data and appends the new data to the end and then applies bit-packing and Zstandard compression and writes them back to the index on disk.

### MetaProFi FASTA/FASTQ indexing

MetaProFi uses the pyfastx^[12]^ python library for parsing FASTA/FASTQ (compressed and uncompressed) files efficiently. However, for MetaProFi’s sequence level indexing, a dedicated input file index is required to be created and shared between multiple processes for fast random access to the sequences. Since sqlite3 that is used in pyfastx to store the index, cannot be shared between multiple processes such that all these processes can be active simultaneously, this can diminish the performance of MetaProFi. So, inspired by the pyfastx python library for indexing FASTA/FASTQ files, we created LMDB (Lightning Memory-Mapped Database) based indexing for FASTA/FASTQ (compressed and uncompressed) files. We used LMDB for its lightning-fast query speed in addition to the capability of sharing the database with multiple processes for querying. In our LMDB database, we build and store a gzip index for the Gzip compressed input files and then store a start offset of every sequence to enable faster random access to any sequence. The FASTA/FASTQ index contains six columns storing the sequence number, name of the sequence, sequence start offset, byte length of the sequence and the number of bases in the sequence. The sequence start offset is used to seek the start of the sequence and the byte length allows us to read the entire sequence.

This minimalistic index is different from what pyfastx builds and this index can be shared between multiple processes for querying random sequences. Further, the index requires less storage compared to pyfstx’s index as we do not store additional information, such as sequence type and read parameters, and also every row in our index is serialized using MessagePack algorithm and then compressed using Zstandard compression algorithm for reducing the storage requirements.

### MetaProFi querying/searching the index

MetaProFi accepts both sequence and FASTA/FASTQ (compressed and uncompressed) files for querying the index. When a multi-sequence file is used for querying, MetaProFi constructs a small LMDB index of the file for distributing the query sequences to multiple cores/processes. MetaProFi extracts *k*-mers and applies hash functions and then in each distributed set MetaProFi constructs a local set data structure using the hash values thus only unique row/bit-slices will be extracted from the index without consuming much memory and time. MetaProFi offers both exact and approximate sequence/*k*-mer queries. Approximate search can be invoked by passing a threshold parameter to MetaProFi’s command-line interface during the querying. In addition to the normal querying of amino acid sequence against the amino acid index and nucleotide sequence against nucleotide index MetaProFi allows querying the amino acid index using nucleotide sequences (metagenomic reads, contigs or assembled genomes). MetaProFi performs a six-frame translation of the nucleotide sequences internally and uses all the six-frames as the query sequence to search the index.

### MetaProFi computing setup

MetaProFi’s performance evaluations were done on a Dell server with the following configuration: AMD EPYC 7702 2.0 GHz CPU with 1.5 TB RAM, Intel SSD DC P4610 3.2 TB (2.9 TiB) and CentOS 7 (kernel 3.10.0-1160.24.1.el7.x86_64) operating system. MetaProFi was installed as a conda environment using Python version 3.7.7 and was allocated with 64 cores, and 60 GiB RAM for all its experiments. All input files were stored on a non-RAID NVMe NFS file system and MetaProFi outputs were stored on the Intel SSD DC P4610 3.2 TB disk.

### MetaProFi indexing of UniProtKB

Two types of MetaProFi index (one at the organism level and one at the sequence level) were constructed for the bacterial sequences in the UniProtKB (Swiss-Prot and TrEMBL) database downloaded in July 2021. The entire Swiss-Prot and TrEMBL datasets were downloaded from UniProt’s FTP site and the accession ids for the bacterial sequences were downloaded by performing a search with “taxonomy:bacteria” on UniProt’s search interface. Three parameters define the architecture of Bloom filters: *m, h*, and *k*, where *m* is Bloom filter size, *h* is the number of hash functions to be applied on every *k*-mer and *k* is the size of the *k*-mer. In MetaProFi, for organism-level indexing, we used the following parameters: *m* = 600,000,000; *h* = 2; *k* = 11. With these parameters, the false positives rate was 0.47, while the false positive rate per query is 10^−6^ if the query sequence size is of a minimum of 35 characters. For sequence-level indexing of the UniProtKB bacterial dataset, we used the same Bloom filter parameters with one change, *m* = 600,000 and with these parameters, the false positive rate was 0.0156 and the per query false positive rate is 10^−6^ if the query sequence size is of a minimum of 34 characters.

### MetaProFi indexing of Tara Oceans dataset

MetaProFi is mainly developed to fill the technology gap of amino acid sequence indexing, but for benchmarking purposes, nucleotide sequence indexing support is also added. With this feature, we downloaded 4 TiB of compressed Tara Oceans dataset (study accession: PRJEB1787) from the ENA archive consisting of 249 samples containing 495 FASTQ files. For MetaProFi indexing of the Tara Oceans dataset, we used the following Bloom filter parameters: *m* = 40,000,000,000; *h* = 1; *k* = 31 and with these parameters the false positives rate was 0.3782 and the false positive rate per query *k*-mer is 10^−6^ if the query sequence size is of a minimum of 50 characters.

### Kmtricks Bloom filter construction for Tara Oceans dataset

Two sets of parameters were used 1) all default values except for the following: *k*-mer size: 31, cores: 64, and Bloom filter size: 40,000,000,000, and 2) same values as 1 but instead of default values for the rest of the command-line parameters, we used the values reported in kmtricks for their Bloom filter construction of the Tara Oceans dataset. Refer data availability section for the full command-line options we used.

### MetaProFi indexing of human RNA-seq dataset

2585 human RNA-seq samples (2.7 TiB) were downloaded from the SRA using the parallel-fastq-dump tool (https://github.com/rvalieris/parallel-fastq-dump), accession numbers obtained from^[5]^. For MetaProFi indexing of this dataset, we used the following Bloom filter parameters *m* = 2,000,000,000; *h* = 1; *k* = 21 taken from^[7]^. For querying we downloaded a fasta file comprising 70,866 transcripts, following^[7]^. We then randomly extracted the first 1000 transcripts using pyfastx^[12]^ and used it for querying the MetaProFi index. RAM utilization was monitored through the Linux command-line utility atop (via the command *atop -mp*).

## Software availability

An open-source implementation of MetaProFi can be found at https://github.com/kalininalab/metaprofi.

## Data availability

In the benchmarks folder (https://github.com/kalininalab/metaprofi/tree/main/benchmarks) of our repository, we provide detailed information on all the commands used for each of the benchmarks and the dataset indexing along with the necessary information to download the data and reproduce our results.

## Acknowledgements

We would like to thank Prof. Sven Rahmann for his helpful suggestions.

## Author contributions

O.V.K. and R.B. conceived the study. O.V.K and S.K.S designed the project, S.K.S designed and implemented the method, developed the software and performed all analyses. S.K helped with the optimization of the performance critical parts of the software. F.D. assisted with pipelines for data analysis and discussion of the results. S.K.S and O.V.K wrote the manuscript, F.D. and S.K. gave detailed feedback on the manuscript. All authors read and approved the manuscript.

S.K.S was partially supported by the UdS-HIPS-Tandem Interdisciplinary Graduate School for Drug Research. S.K was supported by the IMPRS-CS graduate student fellowship and DFG (Deutsche Forschungsgemeinschaft) project number 430158625. O.V.K was funded by the Klaus Faber Foundation.

